# Dysregulation of the SARA–Smurf2 Regulatory Axis in Temporal Lobe Epilepsy

**DOI:** 10.64898/2026.08.10.743913

**Authors:** Estefanía Clavenzani, Juan Manuel Bourbotte Asensio, Laura Ester Montroull, Jesica Piovano, Soledad De Olmos, Marisa Gigena, Sebastián Manuel Bairo, Mariana Bollo, Ariel Martínez, Juan Carlos De Battista, Marco Lisicki, Cecilia Conde

**Affiliations:** Instituto Mercedes y Martín Ferreyra (INIMEC-CONICET-UNC), 5016, Córdoba, Argentina; Hospital de Niños de la Santísima Trinidad de Córdoba, Argentina; Hospital Privado Universitario de Córdoba, Argentina; Facultad de Ciencias Médicas, Universidad Nacional de Córdoba, Córdoba, Argentina; Instituto Universitario Ciencias Biomédicas de Córdoba (IUCBC), X5016, Córdoba, Argentina; Universidad Siglo 21, Córdoba, Argentina

**Keywords:** SARA, Smurf2, Losartan, TGFβ

## Abstract

Temporal lobe epilepsy (TLE) is associated with dysregulation of transforming growth factor β (TGFβ) signaling, a key contributor to epileptogenesis. SARA (Smad Anchor for Receptor Activation), a central regulator of this pathway, is controlled by the E3 ubiquitin ligase Smurf2 through ubiquitination. However, the role of the SARA–Smurf2 axis in regulating TGFβ signaling during TLE has not previously been described, and whether this pathway can be therapeutically targeted remains unknown. Using a pilocarpine-induced status epilepticus (SE) model and astrocytes derived from patients with refractory TLE, we identified dysregulation of the SARA–Smurf2 pathway in both experimental systems. In SE rats, SARA and Glial Fibrillary Acidic Protein (GFAP) levels were significantly increased, whereas Smurf2 induction was insufficient to prevent SARA accumulation. In TLE-derived astrocytes, increased SARA and GFAP immunoreactivity was accompanied by reduced Smurf2 immunoreactivity and altered Smurf2 subcellular distribution. Losartan treatment restored SARA and Smurf2 immunoreactivity toward a control-like pattern in both models and reduced seizure frequency and duration in SE animals. These findings point towards a dysregulation of the SARA–Smurf2 axis as a molecular signature of TLE, support SARA as a potential therapeutic target, providing experimental evidence for the repositioning of Losartan as a potential treatment alternative for drug-resistant epilepsy, warranting further translational and clinical investigation.

**KEY POINTS:**

- Dysregulation of the SARA–Smurf2 axis is a molecular signature of experimental and human temporal lobe epilepsy.
- Impaired Smurf2-dependent regulation of SARA may sustain TGFβ signaling, astrocyte reactivity, and epileptogenesis.
- Losartan restores the SARA–Smurf2 axis and reduces seizures, supporting a novel therapeutic strategy for TLE.

## INTRODUCTION

Temporal lobe epilepsy (TLE) remains a major unmet clinical challenge, particularly in drug-resistant patients. Although anticonvulsant drugs often provide symptomatic control, they are frequently associated with significant side effects hampering adherence and do not modify disease progression. Moreover, 30–40% of patients develop drug-resistant epilepsy, for which surgery —often ablative— remains the primary treatment option^1-2^. This strategy is associated with peri- and post-procedural complications and may lead to irreversible long-term sequelae. In addition, it requires considerable healthcare resources, and its availability is limited to highly specialized centers staffed by specially trained professionals, thereby limiting access for many patients who may require this intervention^3-4^. Therefore, deciphering the molecular mechanisms underlying the development of chronic seizures remains crucial, particularly if these can be reversed using well-tolerated, cost-effective, universally available approaches.

Transforming growth factor β (TGFβ) signaling plays a critical role in epileptogenesis^5^. Smad Anchor for Receptor Activation (SARA), a key adaptor of the TGFβ pathway, regulates TGFβ signaling during neuronal development^6^. Notably, increased expression of SARA, phosphorylated Smad3 (pSmad3), and TGFβ receptor type I (TβRI) has been observed in the hippocampus and cerebral cortex of rats subjected to pilocarpine-induced status epilepticus (SE), as well as in patients with TLE, suggesting dysregulation of this pathway in epileptic tissue^7-8^.

The ubiquitin–proteasome system (UPS) also plays a central role in regulating TGFβ signaling, and its impairment has been proposed as a common pathological feature in several neurological disorders, including epilepsy^9-10^. Smurf2 (Smad Ubiquitination Regulatory Factor 2), an E3 ubiquitin ligase involved in proteasomal degradation, has been shown to target both SARA and TβRs in cell lines^11^.

We have previously demonstrated in the central nervous system that, within the TGFβ pathway, SARA acts as both a positive regulator (promoting Smad2/3 phosphorylation) and a negative regulator (facilitating TβRI dephosphorylation)^6^. Based on these observations, we hypothesized that, in the context of epileptic seizures, increased SARA levels may sustain TβRI activation (phosphorylation), thereby triggering the expression of multiple genes that promote neuronal activity and a pro-inflammatory environment, both hallmark features of this pathology. We investigated whether Smurf2 regulates SARA steady-state levels and whether disruption of this regulatory axis contributes to aberrant TGFβ signaling in epilepsy. (Fig 1A). To this end, we used the rat SE model and astrocytes from human brain tissue of TLE patients. Our results show that Smurf2 dysregulation leads to pathological accumulation of SARA, thereby promoting sustained activation of TGFβ signaling in astrocytes and contributing to epileptogenesis. In pre-clinical models, the efficacy of Losartan [angiotensin II type 1 receptor (AT1) antagonist] in achieving adequate seizure control in TLE^12^ and following SE^13^ remains controversial. However, Zou et al. (2022) showed that in renal interstitial fibrosis, Losartan suppresses TGFβ/Smad signaling by upregulating Smad7, Smurf1, and Smurf2 and downregulating TGFβ, TβRI, TβRII, and phosphorylated Smad2/3^14^. Consistent with the therapeutic potential of this pathway, angiotensin receptor blocker therapy has also been associated with a reduced incidence of epilepsy in humans^15^. Our findings demonstrate that modulation of the TGFβ pathway is associated with the restoration of physiological SARA levels in TLE via Smurfs-TGF-β/Smad pathway, thereby identifying SARA as a potential therapeutic target for the development of novel therapies and providing new mechanistic insight into how Losartan exerts its beneficial effects.

**Figure 1.**
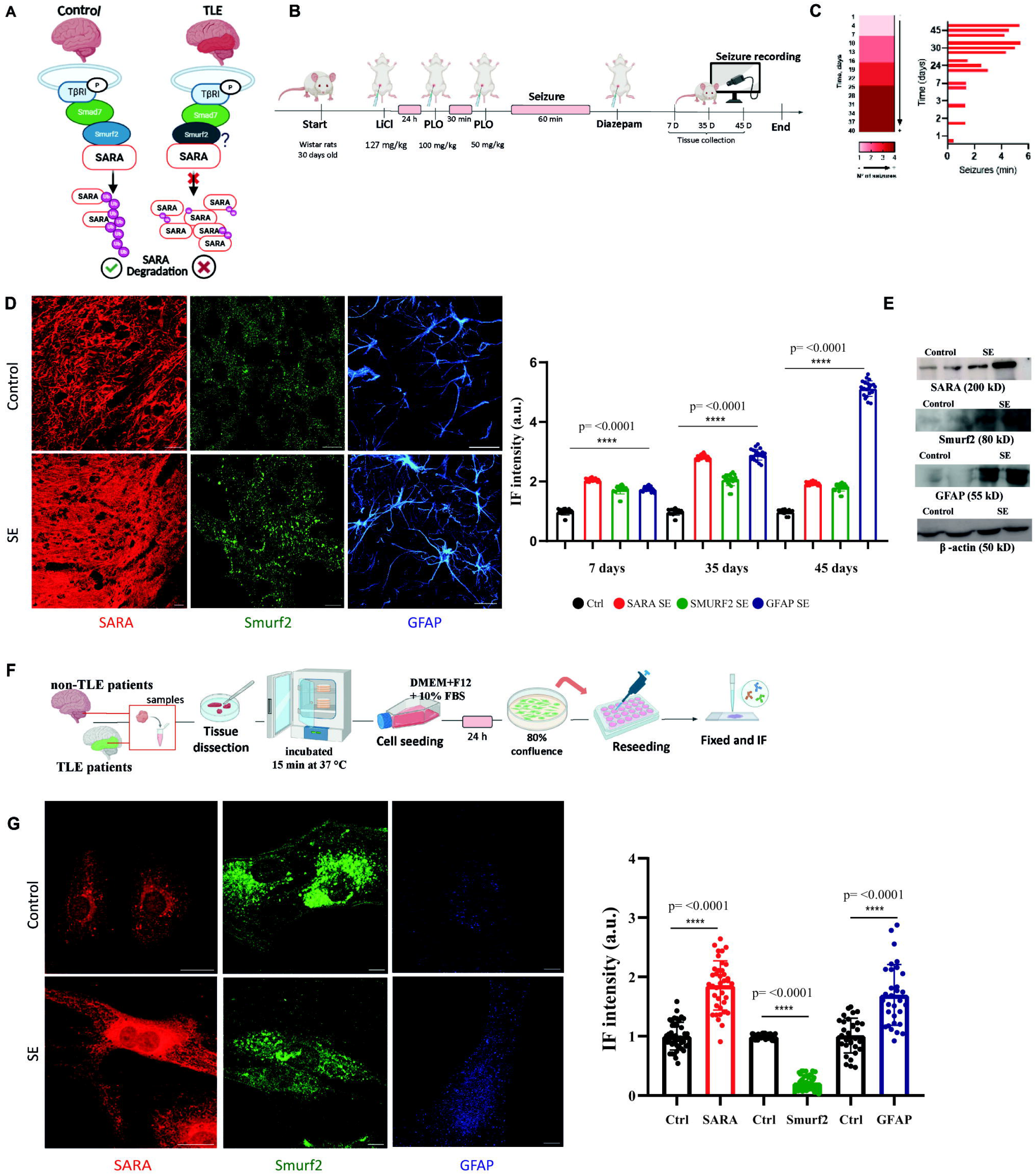
**A. Schematic representation of the regulatory pathway under physiological conditions and the proposed alterations in animal models of SE and in patients with TLE.** Loss of Smurf2-mediated ubiquitination activity is associated with SARA accumulation. Restoring this pathway may represent a potential therapeutic strategy for TLE. **B. Experimental design of the lithium–pilocarpine model of TLE**. Wistar rats (30 days old; 200 g; n=12) received an intraperitoneal (i.p.) injection of lithium chloride (LiCl, 127 mg/kg) 24 h before pilocarpine (PLO) administration. Status epilepticus (SE), defined as continuous Racine stage 4–5 seizures lasting ≥30 min, was induced by two sequential i.p. injections of PLO (100 mg/kg, followed by 50 mg/kg, 30 min later), producing generalized seizures corresponding to Racine stages 4–5. Diazepam (e.g., 10 mg/kg, i.p.) was administered 60 min after SE onset to conclude seizure activity. Animals were monitored by video recording, and brain tissue was collected at 7, 35, and 45 days after SE for subsequent analyses. **C. Characterization of spontaneous recurrent seizures following SE**. Seizure activity was continuously monitored in 3 animals from independent experiments over 45 days following SE. Heat map illustrating the daily number of spontaneous recurrent seizures throughout the experimental period. Daily seizure duration (min) recorded at each time point. **D. SARA, Smurf2 and GFAP expression in hippocampal tissue of rats subjected to SE and controls**. Following transcardial perfusion, 50-μm coronal brain sections were obtained and processed for IF detection of SARA (sc-74493, Santa Cruz Biotechnology, 1:50), Smurf2 (Cat.702291, Invitrogen; 1:50) and GFAP (ZRB2383, Sigma-Aldrich, 1:100). Scale bar: 30 µm. Representative confocal microscopy images (LSM 800 Zeiss, 63X) of the hippocampal CA3 region from control and SE rats showing immunoreactivity of SARA (red), Smurf2 (green), and GFAP-positive astrocytes (blue). Quantification of fluorescence intensity (arbitrary units, a.u.) was performed across the hippocampus. Thirty representative confocal images per animal were analyzed. Experimental data correspond to 2 independent experiments with 6 animals per group in each experiment. Values are expressed as mean ± SEM. Statistical significance was determined using Student’s t-test; ****p < 0.0001 versus control group. **E. Representative Western blot analysis of SARA, Smurf2, and GFAP in cortex of control and SE rats**. Protein extracts from cortex tissue of 2 control and 2 SE rats were analyzed by Western blot using antibodies against SARA (200 kDa), Smurf2 (80 kDa), and GFAP (55 kDa). β-actin (50 kDa) was used as a loading control. The figure shows a representative blot illustrating the relative band intensities observed between experimental groups. **F. Generation of primary human astrocyte cultures from surgical brain tissue**. Cortical tissue samples obtained from pharmacoresistant patients with refractory TLE, and non-epileptic controls were mechanically and enzymatically dissociated using trypsin– EDTA (15 min, 37 °C). The resulting cell suspension was cultured in DMEM/F12 supplemented with 10% FBS until reaching ∼80% confluence, followed by passaging, reseeding, fixation, and IF analysis. **G. Expression of SARA, Smurf2, and GFAP in astrocytes derived from TLE patients and controls**. Representative IF images showing SARA (red), Smurf2 (green), and GFAP (blue) in primary human astrocyte cultures derived from TLE patients and controls. Scale bar: 30 µm. Quantitative analysis of normalized fluorescence intensity (ROI-based measurements) was performed by analyzing 30 astrocytes from each patient in both the TLE and control groups. Data are expressed as mean ± SEM. Statistical significance was assessed using Student’s t-test. ****p < 0.0001. All graphs in this figure were generated using GraphPad Prism 8.

## RESULTS

### SARA increase is associated with Smurf2 dysregulation in an experimental model of Status Epilepticus and human astrocytes from temporal lobe epilepsy

Twelve healthy adults male Wistar rats (200–250 g) were randomly divided into two experimental groups: a control group and a LiCl–PLO group. Rats in the LiCl–PLO group were subjected to lithium chloride–pilocarpine (LiCl–PLO)-induced status epilepticus (SE) as previously described^16^. Rats received an initial dose of PLO (100 mg/kg), followed 30 min later by a second dose of 50 mg/kg. Only animals reaching Racine stages 4–5 were included, as this model closely resembles human temporal lobe epilepsy (TLE)^17^. Control rats received phosphate-buffered saline (PBS). Continuous video monitoring was performed for 24 h at 7, 35, and 45 days after SE induction (Figure 1B). SE animals exhibited a progressive increase in both seizure frequency (from 1 to 4 seizures/day) and seizure duration (from 1 to 6 min) over the study period (Figure 1C).

Immunofluorescence (IF) analysis of hippocampal coronal sections (50 μm; Leica CM1850, Leica Biosystems, IL, USA) revealed significantly increased SARA and GFAP immunoreactivity in the CA3 region of SE rats at 7, 35, and 45 days post-SE compared with controls. SARA immunoreactivity was detected in GFAP-positive reactive astrocytes. Although Smurf2 expression also increased significantly in SE animals (showing approximately 1.6-, 1.8-, and 2.0-fold increases over controls at 7, 35, and 45 days post-SE, respectively), its induction was substantially lower than that of SARA, which increased by approximately 2.2-, 3.0-, and nearly 5.0-fold at the same times (Figure 1D). Western blot analysis confirmed these findings (Figure 1E). These results suggest that the increase in Smurf2 is insufficient to counteract SARA accumulation in SE animals.

We next analyzed SARA, Smurf2, and GFAP by IF in cultured astrocytes derived from surgical specimens of three patients with refractory TLE (Figure 1F). Quantitative analysis revealed significantly increased SARA and GFAP immunoreactivity, together with a marked reduction in Smurf2 immunoreactivity, compared with control astrocytes (Figure 1G).

In addition to the reduction in Smurf2 immunoreactivity, TLE-derived astrocytes exhibited a marked alteration in Smurf2 subcellular localization. In control astrocytes, Smurf2 appeared as small, faint fluorescent puncta or vesicles mainly dispersed predominantly within the perinuclear cytoplasm. In contrast, TLE-derived astrocytes exhibited a diffuse cytoplasmic distribution of Smurf2, with residual Smurf2 immunoreactivity decorating the cell periphery, accompanied by a marked reduction in overall fluorescence intensity. This redistribution is consistent with the reactive phenotype of astrocytes exposed to chronic inflammatory signaling. Together, these findings suggest that Smurf2 dysregulation may promote SARA accumulation and contribute to epileptogenic mechanisms in TLE.

### Losartan reduces SARA levels in an experimental model of SE and in human astrocytes derived from temporal lobe epilepsy tissue

SE rats were analyzed following Losartan treatment, as outlined in Figure 2A. Quantification of daily seizure frequency demonstrated a pronounced reduction of spontaneous recurrent seizures in animals treated with Losartan compared with untreated SE animals, with the effect becoming progressively more evident over the 21-day treatment period. A similar trend was observed for daily seizure duration. No seizure activity was detected in either control group (Figure 2B). Twenty healthy adults male Wistar rats were randomly assigned to four experimental groups: Control, Control + Losartan (LOS), Status Epilepticus (SE), and SE + LOS. Immunofluorescence analysis and quantitative assessment revealed a significant reduction in SARA, TβRI, and GFAP protein levels in the SE + LOS group compared with the untreated SE group (Figure 2C).

**Figure 2.**
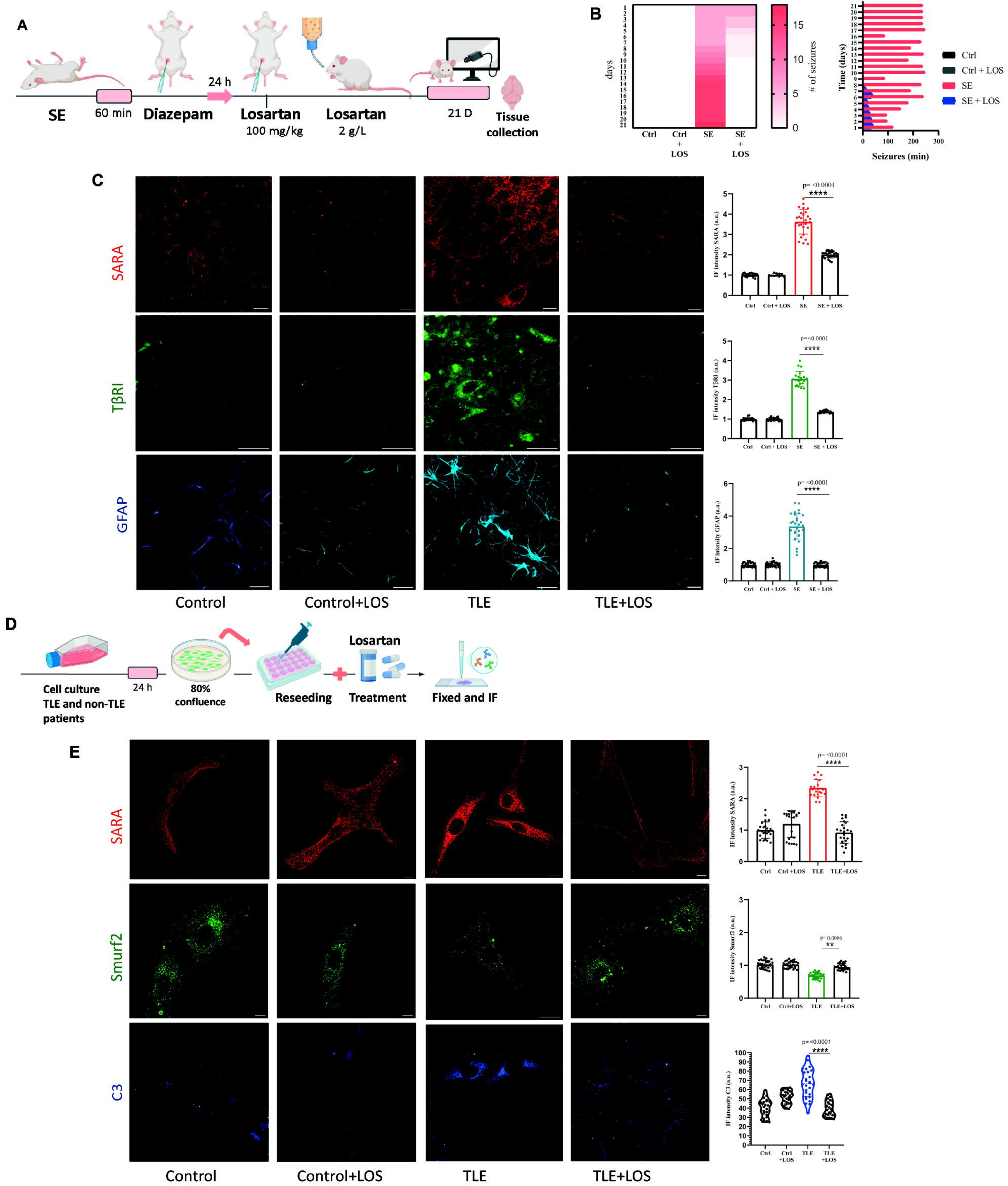
**A. Experimental protocol of SE rats treated with Losartan.** SE rats were generated as outlined in Figure 1A. Animals in the Control + Losartan and SE + Losartan groups received Losartan (100 mg/kg, i.p.), followed by continuous treatment in the drinking water (2 g/L) for 21 days. At the end of the experimental period, animals were perfused, and brain tissue was collected for subsequent analyses. **B. Characterization of spontaneous recurrent seizures and effect of Losartan treatment following SE**. Seizure activity was continuously monitored throughout the 21-day experimental period after SE. Heat map showing the daily number of spontaneous recurrent seizures in the Control (Ctrl), Control + Losartan (Ctrl + LOS), SE, and SE + Losartan (SE + LOS) groups. Daily cumulative seizure duration (min) in each experimental group. **C. Effect of Losartan treatment on SARA, T**β**RI, and GFAP expression in CA3 following SE**. Representative IF images of hippocampal sections from Control (Ctrl), Control + Losartan (Ctrl + LOS), SE, and SE + Losartan (SE + LOS) animals. Scale bar: 30 µm. Immunoreactivity for SARA (red), TβRI (green), and GFAP (blue). Quantification of the IF intensity of SARA, TβRI, and GFAP. Data are expressed as arbitrary units (a.u.) and presented as mean ± SEM. For each experimental condition, 30 images from 3 animals derived from 3 independent experiments were analyzed, n=20. Statistical analysis was performed using Student’s t-test. ****p < 0.0001. **D. Experimental workflow for primary human astrocyte cultures**. Primary astrocytes obtained from TLE patients and non-TLE controls were reseeded onto coverslips and treated with Losartan (1, 2.5, 5, 10, or 20 μM) for 24 or 48 h. Cells were subsequently fixed with 4% paraformaldehyde and processed for IF analysis. **E. Losartan modulates SARA, Smurf2 and C3 expression in primary human astrocytes derived from TLE patients**. Representative IF images of primary astrocytes obtained from 3 controls (Ctrl) and 3 TLE patients, either untreated or treated with Losartan. SARA (red), Smurf2 (green), and C3 (blue). Scale bar: 30 µm. Quantification of IF intensity of SARA, Smurf2, and C3. Data are expressed as mean ± SEM. Statistical significance was determined using Student’s t-test; significance levels are indicated in the graphs. All graphs in this figure were generated using GraphPad Prism 8.

Finally, we analyzed SARA, Smurf2, and complement component 3 (C3, a hallmark of toxic reactive astrocytes recently implicated in the maintenance of neuroinflammation^18^ and epileptogenesis^19^) by IF in human TLE-derived astrocytes following Losartan treatment. Recent evidence indicates that astrocytes are the primary source of C3 in the epileptic hippocampus and that astrocytic C3 expression is progressively upregulated during epileptogenesis^20^. The experimental workflow is shown in Figure 2D. Immunofluorescence images and quantitative analysis revealed a significant reduction in SARA immunoreactivity in Losartan-treated astrocytes. Moreover, Losartan restored both the levels and the subcellular distribution of Smurf2 immunoreactivity to a pattern resembling that observed in control astrocytes. In addition, Losartan significantly reduced C3 immunoreactivity, suggesting an attenuation of the neurotoxic reactive astrocyte phenotype (Figure 2E).

## DISCUSSION

The present study identifies the TGF-β signaling regulators, SARA and Smurf2, as previously unrecognized mediators involved in TLE. Using both a pilocarpine-induced status epilepticus model and astrocytes derived from patients with TLE, we demonstrate a close concordance between the two experimental systems, supporting the translational relevance of our findings. Although substantial progress has been made in understanding TLE pathophysiology—including blood–brain barrier dysfunction, albumin extravasation, neuroinflammation, and activation of the TGFβ signaling pathway—the downstream molecular regulators mediating these processes remain incompletely characterized. Our findings identify SARA and its E3 ubiquitin ligase Smurf2 as novel regulatory nodes within the TGFβ pathway that are altered in epilepsy and, importantly, can be pharmacologically modulated.

Our previous studies demonstrated that SARA accumulation secondary to sustained TβRI activation promotes enhanced TGFβ signaling and synaptic remodeling^6^, processes associated during neuroinflammatory responses and epilepsy. The present findings extend this model by identifying Smurf2 dysregulation as a potential mechanism underlying SARA accumulation in epilepsy. The concomitant increase in SARA together with reduced—or relatively insufficiently increased—Smurf2 levels observed in both experimental systems suggests that impaired Smurf2-dependent regulation of SARA proteostasis may sustain pathological TGFβ signaling, thereby promoting neurotoxic astrocyte reactivity, persistent neuroinflammatory signaling, and ultimately contributing to epileptogenesis. Importantly, Losartan, an angiotensin receptor blocker, restored SARA and Smurf2 immunoreactivity toward a control-like pattern while simultaneously reducing seizure frequency and duration, supporting a functional link between normalization of the SARA–Smurf2 axis and seizure control. These observations provide the first evidence that this regulatory pathway can be pharmacologically modulated in experimental and human TLE.

Although additional studies are required to define the molecular mechanisms linking these events, our data identify the SARA–Smurf2 axis as a novel mechanistic framework underlying persistent TGFβ activation in TLE. Because SARA occupies a downstream position in the TGFβ signaling cascade and has a more restricted biological role than TGFβ itself, restoring Smurf2-dependent control of SARA homeostasis may represent a more selective therapeutic strategy than global inhibition of TGFβ signaling. Together, these findings position the SARA–Smurf2 axis as both a mechanistic framework for understanding sustained TGFβ signaling in epilepsy and a promising therapeutic target for drug-resistant TLE, providing a strong rationale for future translational studies and clinical evaluation. Because Losartan is already an approved drug for hypertension, with a well-established safety profile, the present findings may facilitate the rapid translation of SARA-targeted therapeutic strategies into clinical studies for drug-resistant TLE.

## AUTHOR CONTRIBUTIONS

E.C. performed the experiments, analyzed the data, and contributed to writing the manuscript. J.M.B.A. assisted with immunofluorescence (IF) experiments and image analysis. L.E.M. and S.D.O. assisted with animal perfusion, Western blot (WB) experiments, and data analysis. J.P. and M.G. assisted with animal handling. S.B. and M.B. contributed to astrocyte culture procedures. J.C.D.B. provided surgical tissue samples and contributed to the discussion. A.M. contributed to the interpretation and discussion of the results. M.L. contributed to the study design and interpretation of the results. C.C. conceived the study, designed the experiments, supervised the research, and drafted the manuscript. All authors reviewed and approved the final version of the manuscript.

## ACKNOWLEDGMENTS

We thank Dr. Andrea Pellegrini for assistance with the cell culture facility and Eliana Martínez for technical collaboration in animal handling. We are grateful to Dr. Mauricio Martin for his support and comments during the experiments.

## REFERENCES

1. Asadi-Pooya AA, Stewart GR, Abrams DJ, Sharan A. Prevalence and Incidence of Drug-Resistant Mesial Temporal Lobe Epilepsy in the United States. World Neurosurg. 2017; 99:662–666.

2. Perucca E, Perucca P, White HS, Wirrell EC. Drug resistance in epilepsy. Lancet Neurol. 2023; 22(8):723–734.

3. Jehi L, Friedman D, Carlson C, Cascino G, Dewar S, Elger C, Engel J Jr, Knowlton R, Kuzniecky R, McIntosh A, O’Brien TJ, Spencer D, Sperling MR, Worrell G, Bingaman B, Gonzalez-Martinez J, Doyle W, French J. The evolution of epilepsy surgery between 1991 and 2011 in nine major epilepsy centers across the United States, Germany, and Australia. Epilepsia. 2015; 56(10):1526–33.

4. Bartolomei F, Makhalova J, Benoit J, Lagarde S. The different subtypes of temporal lobe seizures networks. Rev Neurol (Paris). 2025; 181(5):368–381.

5. Luo J. TGF-β as a Key Modulator of Astrocyte Reactivity: Disease Relevance and Therapeutic Implications. Biomedicines. 2022; 10(5):1206.

6. Rozés-Salvador V, Wilson C, Olmos C, Gonzalez-Billault C, Conde C. Fine-Tuning the TGFβ Signaling Pathway by SARA During Neuronal Development. Front Cell Dev Biol. 2020; 8:550267.

7. Yu W, Du Y, Zou Y, Wang X, Stephani U, Lü Y. Smad anchor for receptor activation contributes to seizures in temporal lobe epilepsy. Synapse. 2017; 71(3).

8. Zhang W, Du Y, Zou Y, Luo J, Lü Y, Yu W. Smad Anchor for Receptor Activation and Phospho-Smad3 Were Upregulated in Patients with Temporal Lobe Epilepsy. J Mol Neurosci. 2019; 68(1):91–98.

9. Caldeira MV, Curcio M, Leal G, Salazar IL, Mele M, Santos AR, Melo CV, Pereira P, Canzoniero LM, Duarte CB. Excitotoxic stimulation downregulates the ubiquitin-proteasome system through activation of NMDA receptors in cultured hippocampal neurons. Biochim Biophys Acta. 2013; 1832(1):263–74.

10. Caldeira MV, Salazar IL, Curcio M, Canzoniero LM, Duarte CB. Role of the ubiquitin-proteasome system in brain ischemia: friend or foe? Prog Neurobiol. 2014; 112:50–69.

11. Wojtowicz S, Lee S, Chan E, Ng E, Campbell CI, Di Guglielmo GM. SMURF2 and SMAD7 induce SARA degradation via the proteasome. Cell Signal. 2020; 72:109627.

12. Reyes-Garcia SZ, Scorza CA, Ortiz-Villatoro NN, Cavalheiro EA. Losartan fails to suppress epileptiform activity in brain slices from resected tissues of patients with drug-resistant epilepsy. J Neurol Sci. 2019; 397:169–171.

13. Nozaki T, Ura H, Takumi I, Kobayashi S, Maru E, Morita A. The angiotensin II type I receptor antagonist Losartan retards amygdala kindling-induced epileptogenesis. Brain Res. 2018; 1694:121–128.

14. Zou J, Zhou X, Ma Y, Yu R. Losartan ameliorates renal interstitial fibrosis through metabolic pathway and Smurfs-TGF-β/Smad. Biomed Pharmacother. 2022; 149:112931.

15. Liu J, Liu Q, Jiang X, Hu Y, Xu J, Li Y, Ma M, Fang J, Zhou D, He L. Angiotensin-converting enzyme inhibitors and angiotensin-receptor blockers usage is associated with a reduced risk of seizures and epilepsy after ischemic stroke. Neurol Sci. 2026; 47(3):311.

16. Du Y, Zou Y, Yu W, Shi R, Zhang M, Yang W, Duan J, Deng Y, Wang X, Lü Y. Expression pattern of sorting Nexin 25 in temporal lobe epilepsy: a study on patients and pilocarpine-induced rats. Brain Res. 2013; 1509:79–85.

17. Inoue O, Sugiyama E, Hasebe N, Tsuchiya N, Hosoi R, Yamaguchi M, Abe K, Gee A. Methyl ethyl ketone blocks status epilepticus induced by lithium-pilocarpine in rats. Br J Pharmacol. 2009; 158(3):872–8.

18. Lian H, Yang L, Cole A, Sun L, Chiang ACA, Fowler SW, Shim DJ, Rodriguez-Rivera J, Taglialatela G, Jankowsky JL, Lu HC, Zheng H. NFκB-Activated Astroglial Release of Complement C3 Compromises Neuronal Morphology and Function Associated with Alzheimer’s Disease. Neuron. 2015; 85(1):101–115.

19. Liddelow SA, Olsen ML, Sofroniew MV. Reactive Astrocytes and Emerging Roles in Central Nervous System Disorders. Cold Spring Harb Perspect Biol. 2024; 16(5):a041356.

20. Jiang M, Romagnolo A, Aronica E, Wang Y. Dynamic astrocytic complement C3 activation in the epileptic hippocampus. Front Neurol. 2025; 16:1682488.

